# Activation of Heme Metabolism Promotes Tissue Health After Intraarticular Injury or Surgical Exposure

**DOI:** 10.1101/2024.05.29.596521

**Authors:** Suryamin Liman, Piedad C. Gomez-Contreras, Madeline R. Hines, Emily Witt, Jacob S. Fisher, Kevin J. Lu, Lauren D. McNally, Alicia T. Cotoia, Maxwell Y. Sakyi, Brett A. Wagner, Michael S. Tift, Jessica E. Goetz, James D. Byrne, Mitchell C. Coleman

## Abstract

Posttraumatic osteoarthritis (PTOA) is a well-recognized public health burden without any disease modifying treatment. This occurs despite noted advances in surgical care in the past 50 years. Mitochondrial oxidative damage pathways initiate PTOA after severe injuries like intraarticular fracture that often require surgery and contribute to PTOA after less severe injuries that may or may not require surgery like meniscal injuries. When considering the mitochondrial and redox environment of the injured joint, we hypothesized that activation of heme metabolism, previously associated with healing in many settings, would cause prototypic mitochondrial reprogramming effects in cartilage ideally suited for use at the time of injury repair. Activation of heme metabolism can be accomplished through the gasotransmitter carbon monoxide (CO), which activates hemeoxygenase-1 (HO1) and subsequent heme metabolism. In this study, we employed unique carbon monoxide (CO)-containing foam (COF) to stimulate heme metabolism and restore chondrocyte oxygen metabolism *in vitro* and *in vivo*. Doxycycline-inducible, chondrocyte-specific HO1 overexpressing transgenic mice show similar mitochondrial reprogramming after induction compared to COF. CO is retained at least 24 h after COF injection into stifle joints and induces sustained increases in heme metabolism. Lastly, intraarticular injection of COF causes key redox outcomes without any adverse safety outcomes in rabbit stifle joints *ex vivo* and *in vivo*. We propose that activation of heme metabolism is an ideal adjuvant to trauma care that replenishes chondrocyte mitochondrial metabolism and restores redox homeostasis.

## [Introduction]

Heme metabolism plays a key role in restoring the redox, mitochondrial, and energetic pathways of mammalian cells after injuries or during the resolution of inflammation [1-7]. Stimulating heme metabolism with prior generations of CO donors [8-10] or other means of upregulating the initiating enzyme, hemeoxygenase-1 (HO1) has shown how this pathway provides some degree of protection against inflammatory models of arthritis [11-14] and a wide variety of non-orthopedic pathologies [15-18]. Many of these studies have concentrated on the anti-inflammatory features of these agents, but CO and heme metabolism are also known to improve mitochondrial function [7, 19]. This is observed throughout the body despite its well-known role in inhibiting respiration at high doses. CO appears to upregulate mitochondrial electron transport chain (ETC) activity and antioxidant pathways, but relationships between heme metabolism, mitochondria, and redox status in healthy, primary, articular chondrocytes have not been described. Our research group has focused on early features of posttraumatic osteoarthritis (PTOA) in articular chondrocytes, showing how mitochondrial damage and dysfunction associated with traumatic injury can initiate PTOA. As mitochondrial reprogramming [20, 21] approaches are increasingly developed and applied, we suggest that heme metabolism’s coordination of chondrocyte redox and mitochondrial physiology may provide prototypic strategies for cartilage protection and restoration.

Our group and others [22, 23] have shown how sublethal mechanical injuries to articular cartilage initiate oxidative damage to the mitochondria over the first few hours that can be prevented by augmenting intracellular glutathione (GSH) or inhibiting mitochondrial electron transport [24, 25]. This results in mitochondrial dysfunction that does not resolve on its own but goes on to contribute to PTOA in multiple preclinical models [26]. Prior studies have also identified damage from intraarticular surgical exposures including air [27-29], low osmolarity [30, 31], unbuffered pH [32], and iatrogenic injuries [33]. When considering augmentation of the preventive opportunity offered by repair of traumatic injuries, we hypothesized that independent of mechanical joint injury, damage from these other sources associated with surgery might mitigate some of the benefits of surgery. We propose this could occur *via* damage to chondrocyte mitochondria, rendering them dysfunctional and unable to support healthy cellular metabolism. Thus, the effects of joint injuries on chondrocyte mitochondria might be compounded by subsequent repairs that expose intraarticular tissue to air or irrigations with low osmolarity, unbuffered surgical saline. We further hypothesized that activation of heme metabolism with acute delivery of CO to damaged or exposed articular cartilage might mitigate the effects of each of these injuries, improve mitochondrial metabolism, and prevent oxidative damage and stress.

To test this hypothesis, we applied the CO-containing foam (COF) developed by Byrne *et al.* [16] to tissue culture models, primary bovine osteochondral explants, and lapine and murine stifle joints. To better examine chondrocyte mitochondrial reprogramming by heme metabolism as a prototypic mitochondrial reprogramming, this was complemented with genetic overexpression of HO1 using the TetOn inducible system under a chondrocyte-specific promoter. We observed consistent increases in mitochondrial function and ETC enzyme complement with COF that were associated with increased activity and decreased oxidation of the GSH pathway. These effects were observed in the presence and absence of traumatic injury and were recreated in HO1 overexpressing chondrocytes *in vivo*. COF proved to be well tolerated, demonstrating no toxicity in any species tested. This exciting, novel material or related mitochondrial reprogramming approaches may prove a disease-modifying adjuvant to surgical care, augmenting the benefits of orthopedic surgical care in any setting where orthopedic trauma, iatrogenic cartilage injury, or joint exposure is a patient concern.

## [Materials and Methods]

### Cell and Tissue Culture

Primary bovine chondrocytes extracted from healthy cartilage as well as the human chondrocyte cell line T/C-28a2 (Millipore, Temecula, CA) were used for monolayer studies. Primary chondrocytes were extracted via digestion and centrifugation. Briefly, cartilage from the explants were digested with collagenase and pronase (0.1 mg/ml; Sigma-Aldrich) overnight. The chondrocytes were then collected after pelleting, resuspended, and plated.

Cylindrical bovine osteochondral explants were utilized for injury studies. Intact, fresh bovine stifle (knee) joints obtained from a local abattoir (Bud’s Custom Meats, Riverside, IA) were dissected and both medial and lateral femoral condyles were used. Cylindrical explants 10 mm in diameter were obtained from the load bearing region of each femoral condyle. The explants were washed in Hank’s balanced salt solution (HBBS) with 50 U/mL penicillin, 50 µg/mL streptomycin, and 2.5 µm/mL amphotericin B, then placed into culture medium. Both explant and monolayer cultures were maintained in cell culture medium with 45% Dulbecco’s Modified Eagle’s medium, 45% F-12, 10% fetal bovine serum (FBS) (Gibco), 100 U/mL penicillin, 100 µg/mL streptomycin, and 2.5 µm/mL amphotericin B. All specimens were kept in a tissue culture incubator maintained at 5% O_2_, 5% CO_2_ and 37 °C.

### Mechanical Injury

After overnight equilibration to culture conditions, the mechanical impact injury was delivered *via* drop tower as previously described [24]. A 2 J/cm^2^ of impact was applied with a flat, stainless steel impermeable platen. The explants were leveled and adhered to a 3 x 3 cm stainless steel plate with polycaprolactone prior to impact. Release of the drop tower was secured to ensure a single impact. The explants were then rinsed with HBSS before being returned to normal cell culture medium.

### Gas Delivery *via* Gas Entrapping Foam

Novel materials were developed to entrap room air, nitrogen, or CO. The material includes 0.5% wt xanthan gum (Modernist Pantry), 0.8% wt methylcellulose (Modernist Pantry) in phosphate buffered saline (PBS). The facilitate gas entrapment, the materials were pressurized in a custom-made whipping siphon up to 200 PSI of the gas of interest according to Byrne *et al.* [16]. Foams were applied for 30 min for monolayer cell cultures or 1 h for explants.

### *Ex vivo* Stifle Culture

Intact but immediately post-mortem animals from other studies were used to explore acute effects of COF and room air foam (RAF) within the joint. New Zealand White rabbits, female aged 8 to 9 months (Charles River) and C57B6J mice, both sexes, aged 8 to 15 weeks (Jackson) were utilized. After euthanasia, foam was introduced into the stifle joint *via* injection (rabbits with a 27 g needle, mice with a 31 g needle) immediately medial of the patellar tendon and the presence of foam within the joint was confirmed *via* fluoroscopy and inclusion of 20% Isovue-370 in the foam material. Stifles were extended and flexed gently and then left for 30 min. Lapine injections were 0.5 ml foam and murine injections were 0.05 ml foam.

### Measurement of CO after Intraarticular Injection

To assess CO concentrations after intraarticular injection, mice were given injections of 0.05 ml RAF or COF under anesthesia, returned to their cages and allowed to move freely, then stifles were harvested 24 h later and frozen. For analysis, pre-weighed stifles were rinsed free of blood with PBS and placed in 2 mL bead mill tubes with stainless steel beads (2.4 mm) and diluted 10-fold with ice cold water. Samples were then homogenized on a bead-mill homogenizer (VWR), placed in an ultrasonic bath (Branson) for 5 minutes. Using a gas-tight syringe (Hamilton), 20 µl of homogenate was then added through a septum to a CO-free amber borosilicate glass vial that contained 20 µl of sulfosalicylic acid (20%). The CO released into headspace of the vial was then flushed through a gas chromatograph with a reducing compound photometer (Peak Labs; Peak Performer 1) and was quantified using a standard curve created daily using different volumes of 1ppm CO gas (AirGas).

### Assessments of Intraarticular Injection of COF

To investigate potential toxicity from injection of COF, we utilized New Zealand White rabbits, female aged 8 to 9 months (Charles River). We compared stifle joints from rabbits receiving no injection (Normal) to stifles from rabbits the received an injection of RAF in one stifle and COF in the opposing stifle, alternating left and right for either injection in individual animals that were then euthanized 1 week after injection or 4 weeks after injection, n = 10. To assess synovial health or any damage from injection, we stained cross-sections of synovium with hematoxylin and examined synovial thickness using Olympus Desktop (Olympus) and scored cellularity of synovia as the number of cells thick the tissue presented in these cross-sections. To assess chondrocyte viability from these stifles, we stained the tibial articular surfaces with Calcein AM (Life Technologies), imaged *via* confocal microscopy, and counted viable cells per field using ImageJ.

### Western Blot

Protein extracts from cell models, explants, and tissue specimens were denatured and reduced by addition of LDS sample buffer (Invitrogen) and DTT reducing agent (Invitrogen), heated at 70°C by 10 min. A total of 20 ug of protein was loaded per well and electrophoresed through a 10% Bis-Tris NuPage acrylamide gel (Invitrogen), at 120 Volts constant by 2h with MES-SDS running buffer (Invitrogen). After the electrophoresis, the gel proteins were transferred to a 0.2 uM PVDF membrane (BioRad), at 300 mAmp constant by 2h with 20% methanol 0.1% SDS transfer buffer (Invitrogen). The blotted membrane was stained using Ponceau S, as a loading and protein quality control. The membrane was blocked by 30 min with Bovine serum albumin (BSA) 5% solution. The primary antibodies mitofusin-1 (Abcam), sirtuin 1 (Cell Signaling), mitochondrial electron transport chain complexes I-V (cocktail, Abcam), p65 (Cell Signaling), OPA-1 (Cell Signaling) were diluted 1:1000 in BSA 2.5% and incubated overnight at 4°C in a rotator shaker. After TBST rinsing, HRP-linked secondary abs (Cell Signaling) diluted 1:2000 in BSA 2.5% were incubated by 1h RT. Once TBST rinsed, the protein signal was developed by Super Signal West Femto (Thermo) and visualized with Amersham Hyperfilm ECL system (GE Healthcare) film.

### Extracellular Flux Analyses

To evaluate mitochondrial activity by live chondrocytes as rapidly after COF treatment as possible, 20,000 primary bovine chondrocytes per well were plated on XF96 Extracellular Flux Analyzer (Seahorse Bioscience, Agilent) plates. After 3 days to allow attachment, wells were exposed to either RAF or COF then media was changed and standard mitochondrial stress test measures were conducted as previously described [25]. Briefly, the following reagents were successively injected into each well (final concentration in the well shown): oligomycin (2 μM), carbonyl cyanide-4- (trifluoromethoxy)phenylhydrazone (FCCP) (250 nM), antimycin A (5 μM), rotenone (2 μM). After completion of the assay, the number of cells was counted. The oxygen consumption rates (OCR) were then normalized to the number of cells. Basal respiration is reported as basal OCR minus OCR after rotenone and antimycin A injection. ATP-linked respiration is reported as basal OCR minus OCR after oligomycin injection. Proton leakage is OCR after oligomycin injection minus OCR after rotenone and antimycin A injection, normalized to each specimen’s basal respiration.

### ETC Complex Activity Assays

Electron transport chain activities were measured as previously described [34]. Briefly, Complex II samples were incubated with or without succinate in 25 mM potassium phosphate buffer, 5 mM magnesium chloride, 2 mM potassium cyanide, and BSA (2.5 mg/ml) for 10 min at 30°C. After 10 min, antimycin A (200 μg/ml), rotenone (200 μg/ml), 5 mM 2,6-dichloroindophenol (Sigma-Aldrich), and 7.5 mM coenzyme Q1 were added. The activity was calculated with this equation *A* = *ε* · *c* · *l*, where *A* is the absorbance, *ε* is the molar extinction coefficient, *c* is the concentration, and *l* is the path length. The concentration was calculated by dividing difference of the absorption at 600 nm with molar extinction coefficient of 2,6-dichloroindophenol, 19.1, and the length, 1 cm. The number was normalized by the protein content. Complex IV samples were assayed in 25 mM potassium phosphate buffer, 0.5 mM n-dodecyl β-maltoside, and 1.5 mM reduced cytochrome c. The activity was calculated as difference of the absorption at 550 nm, divided by molar extinction coefficient of cytochrome *c*, 19.6, and then normalized to each sample’s protein content.

### Cartilage-Specific HO1 Overexpression and Destabilization of the Medial Meniscus

Cartilage-specific HO1 overexpression was achieved using a doxycycline-inducible overexpression system implemented in two strains of transgenic mice that when crossed, provide HO1 overexpression upon introduction of doxycycline. We created a *Col2A1-TetOn^+/-^* strain to enable the necessary cartilage-specific expression of TetOn, and a Tet response element (TRE)-*mHMOX1*, *TRE-mHMOX1^+/-^* strain to provide doxycycline-responsive HO1 overexpression. Both strains were created *via* knock-in on a B6 background with the help of the Genome Editing Facility at the University of Iowa, which performed all cloning, embryo injection, and genetic characterization of this strain. Briefly, Col2A1-TetOn mice have a knock-in, optimized, second-generation reverse tetracycline transactivator (*rtTA*M2*) gene [35] under a promoter taken from a portion of *Col2A1* [36] that provides chondrocyte specific expression *in utero* [36] up to the 10-15 week age groups shown in this study. Briefly, the inserted *Col2A1-TetOn* transgene contains an SV40 polyadenylation signal, the previously characterized 2 x 182 bp doublet taken from the *Col2A1* promoter sequence [36], a minimal woodchuck hepatitis virus post-transcriptional regulatory element, the TetOn gene [35], a Factor Xa site and a polyadenylation signal. TRE-mHMOX1 mice have a different gene knocked-in containing the murine *HMOX1* under the TRE promoter [35]. Similar to the Col2A1-TetOn mouse, but with distinct promotion and payload, the inserted *TRE-mHMOX1* transgene contains an SV40 polyadenylation signal, the tet response element critical for rtTA*M2 binding (7 repeats of TCCCTATCAGTGATAGAGA separated by spacer sequences), a minimal cytomegalovirus promoter, the Kozak sequence and *mHMOX1*, a β-globin intron and a polyadenylation signal. Double transgenic *Col2A1-TetOn^+/-^, TRE-mHMOX1^+/-^* mice were confirmed with genotyping and both strains are maintained separately as heterozygotes. We have provided supplemental Figures 1 and 2 to show the transgenes that have been inserted. For pivotal studies, mice aged 8-15 weeks were fed between 1-5 days of doxycycline-containing chow (dox chow, TD.01306 at 625 mg/kg, Teklad), with stable overexpression observed after day 1. Stifles for histological studies [37] and specimens for Western blots were prepared as previously described [37].

To explore injuries and the development of PTOA in these mice, we applied the destabilization of the medial meniscus (DMM) model commonly in use [38]. Mice underwent DMM to the right stifle after reaching skeletal maturity (12-16 weeks). DMM was performed by opening the skin and joint capsule along the patellar ligament. After the joint was open the medial meniscal tibial ligament was severed then the joint capsule and skin was sutured closed [38]. Animals were allowed to move freely around their enclosure after the surgery until the time of euthanasia. Both injured and contralateral control stifles were harvested and processed for histology. Degree of PTOA was evaluated using OARSI scoring of the medial compartment.

### Immunofluorescence (IF)

Slides were deparaffinized and rehydrated via a series of washes: 100% xylene, 100% ethanol, 95% ethanol, 80% ethanol, and distilled water. Slides were then placed in 0.1 M sodium citrate, pH 6, overnight, up to 16 h, at 55 °C. The samples were rinsed, and peroxidases were quenched. Samples were blocked using normal goat serum-blocking solution. The primary anti-HO1 antibody was incubated overnight (1:150) at 4 °C. The secondary antibody (1:200) was incubated 30–40 min at room temperature. The samples were developed using fluorescent developing reagent kits utilizing Cy5 channels (Ventana, Roche). Quantum Dot DAPI (Ventana, Roche) was utilized for counterstaining.

### Redox Status Visualization *via* Live Cell Imaging

In order to assess the redox status, cell viability, and mitochondria of chondrocytes *in situ*, we applied live cell stains to cultured tissue and visualized cross sections of the cartilage using confocal microscopy. First, the explants were cut with a precision saw (IsoMet 1000, Buehler) and a clean cross section was revealed *via* removal of a 0.25 mm deep portion of the face of the cartilage with a scalpel. Monochlorobimane (MCB, Invitrogen) was applied to reveal the <u>reduced</u> (as opposed to oxidized) thiols, alongside Calcein AM, to assess cellular viability. MCB reacts with endogenous glutathione-S-transferase and reduced thiols (predominantly GSH) to create a blue fluorescent complex. This blue fluorescence thus indicates the presence of reduced thiols and decreases in that blue staining would indicate oxidation. Both explants and cells were stained in 20 µM MCB and 1 µM calcein AM. All staining and imaging occurred in DMEM/F-12 media without phenol red (Gibco). The samples were then observed with the confocal microscopy with using excitations at 405 nm and 488 nm for MCB and calcein AM, respectively.

For mitochondrial staining, the explants or cells were stained with MitoTracker Deep Red (MTDR) (Invitrogen) at 1 µM and co-stained with 1 µM of calcein AM, for cell viability. The samples were observed with the confocal microscopy with excitation at 633 nm for MTDR and 488 nm for calcein AM. Cartilage images were exported, manually segmented according to appropriate depths or areas, and analyzed with a previously published, custom MATLAB-based algorithm [39]. This algorithm uses calcein AM to identify live cells only and then determines intracellular MCB or MTDR fluorescent intensity in each live cell.

### Biochemical Measurement of GSH and GSSG

Griffith’s GSH/GSSG assay as previously described [40] was deployed to confirm MCB results. Cartilage samples cut from explants were minced into 5% sulfosalicylic acid and subjected to three freeze-thaw cycles to lyse the cells. Tissue and cell lysates were then mixed with buffer solution containing dithionitrobenzoic acid (DTNB), GSH reductase, and NADPH (all from Sigma-Aldrich). GSH reacts with DTNB to form a yellow product, while the GSH reductase recycles the GSH. As a result, the absorbance rate change at 412 nm as measured *via* spectrophotometer is proportional to GSH concentration. All sample concentrations were determined via comparison to appropriate standard curves. For GSSG determination, 20% sample volume of 2-vinylpyridine (Sigma-Aldrich) was added to sample aliquots for 1 h before analysis to quench GSH and prevent the reaction of <u>reduced</u> GSH with DTNB.

### Image Analysis & Statistics Analysis

The MCB or MTDR intensity was quantified using a custom MATLAB-based algorithm [39]. The algorithm first detects live cells using the calcein AM channel and creates regions of each image defined by these cells. It then quantifies the MCB or MTDR intensity within these regions, ensuring the measurements are specific to live cells. Average intensity per cell was obtained and the average intensity of cells were compared in zones and injury/ non-injury group with one-way ANOVA analysis.

## [Results]

### CO Rapidly Induces HO1 and Increases Mitochondrial Function

We first wanted to demonstrate that COF induces HO1 in articular cartilage, as shown in other tissues [41, 42]. Following 30 min exposure to either CO or RAF, cartilage samples were harvested between 0 and 12 h after exposure for western blotting of explant cartilage. Our results showed that CO induces HO1 as soon as 1 h after foam exposure. This elevation of HO1 lasted for at least 12 h after the exposure, **Fig. 1A**. We also observed increases in mitofusin-1, a marker of mitochondrial fusion previously associated with CO exposure as well as mitochondrial health in other tissues. Based on prior reports [43-45], we hypothesized that exogenous CO exposure improves the mitochondrial respiratory function of chondrocytes. We tested this hypothesis using primary chondrocytes extracted from bovine cartilage subjected to mitochondrial stress tests as rapidly after foam treatment as possible. This measurement was approximately 1.5 h after initial exposure, extracellular flux equilibration, and early portions of the mitochondrial stress test. The result showed COF-treated chondrocytes had higher oxygen consumption rates (OCR) in the basal and ATP-linked portions of the mitochondrial stress test, as shown in **Fig. 1B and 1C**, without increasing the percentage of proton leakage, **Fig. 1D**, suggesting no increases in oxidative stress or damage to electron transport chains associated with increased activity. These studies demonstrated that COF increased primary chondrocytes’ HO1 content and augmented their mitochondrial function.

**Fig. 1.**
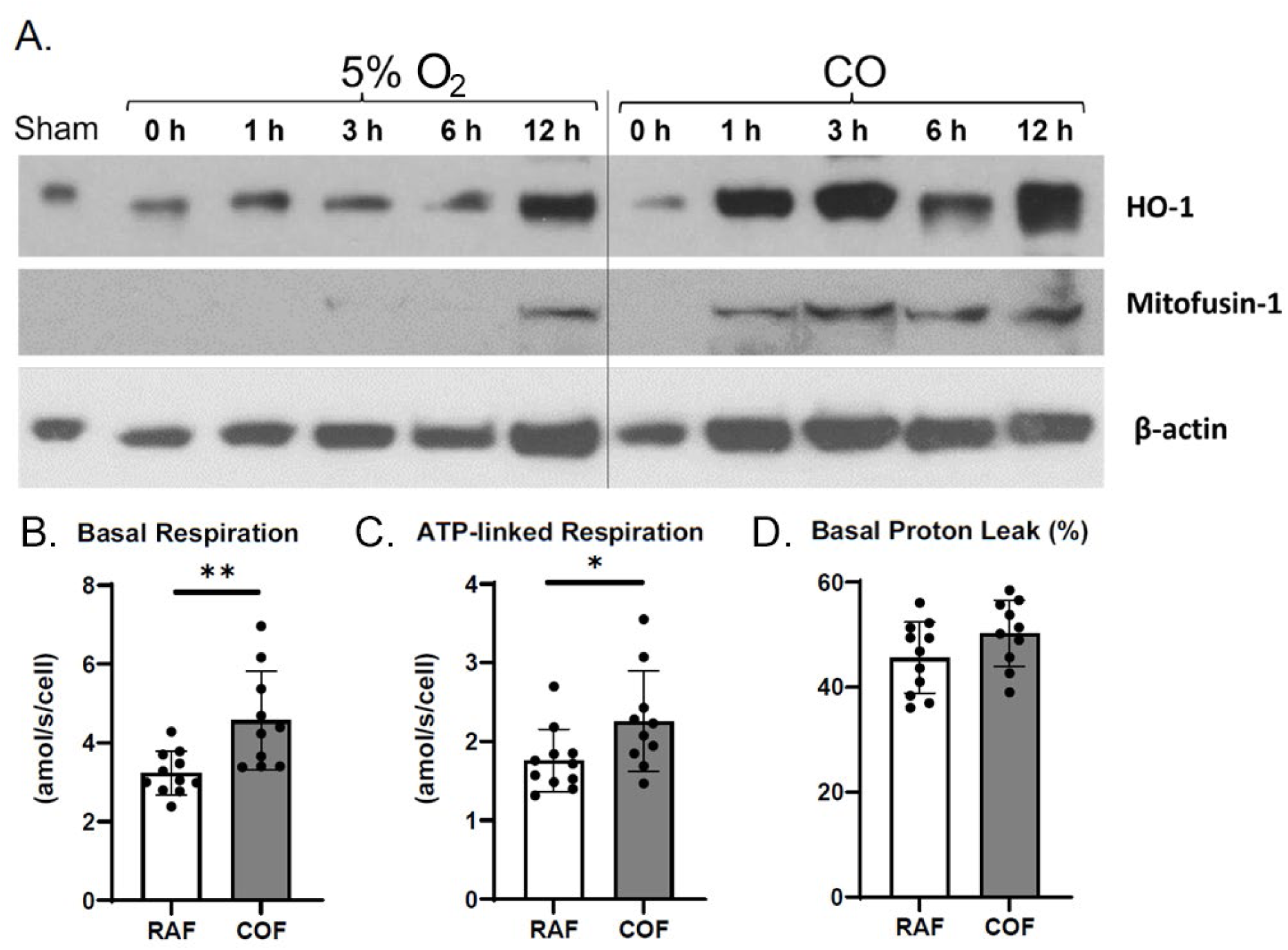
CO Rapidly Induces Chondrocyte HO1 and Increases Mitochondrial Function. Western blots of tissue homogenates from explants at various timepoints after CO exposure show increasing presence of HO1 within 1 h after CO compared to control specimens exposed to normal 5% O2 gassed solutions (A). This cohered with increased presence of mitofusin-1. Mitochondrial stress tests of chondrocytes extracted from similarly treated cartilage showed increased per-cell basal respiration (B), ATP-linked respiration (C), and no indications of damage to the mitochondrial electron transport chain as indicated by a lack of change in proton leakage (D). n = 9, p < 0.05 by t-test.

### HO1 Overexpression in Articular Chondrocytes *In Vivo* Increases Mitochondrial Metabolism and Decreases Basal p65 Expression

Because culturing primary chondrocytes is associated with a variety of well-known mitochondrial artifacts, to better examine the effects of increasing HO1 and activating heme metabolism in primary chondrocytes *in vivo* we designed *Col2A1-TetOn^+/-^, TRE-mHMOX1^+/-^* mice to overexpress HO1 in articular chondrocytes in the presence of doxycycline. We fed mice doxycycline-containing chow for 1-5 days and examined immunofluorescent staining for HO1 and western blots for HO1. Our result showed that by 2 days of dox chow feeding, increased HO1 was seen in cartilage but not in meniscus, **Fig. 2A quantified in 2B**. To examine ETC content with HO1 overexpression, western blots for complexes I, II, III, IV, and V were run on cartilage from murine hind foot preparations, **Fig. 2C**. These samples also showed elevation of the pro-mitochondrial sirtuin 1 (Sirt1) and decreased basal p65 staining after doxycycline feeding. We saw similar changes in Sirt1 with 1 and 3 days of dox chow in both the forelimb paw and hindlimb paw, suggesting that effects on mitochondrial pathways were sustained over these timepoints, **Fig. 2D**. These data demonstrate *in vivo* that the activation of heme metabolism triggers key molecular effects in articular chondrocytes corresponding to the effects of COF.

**Fig. 2.**
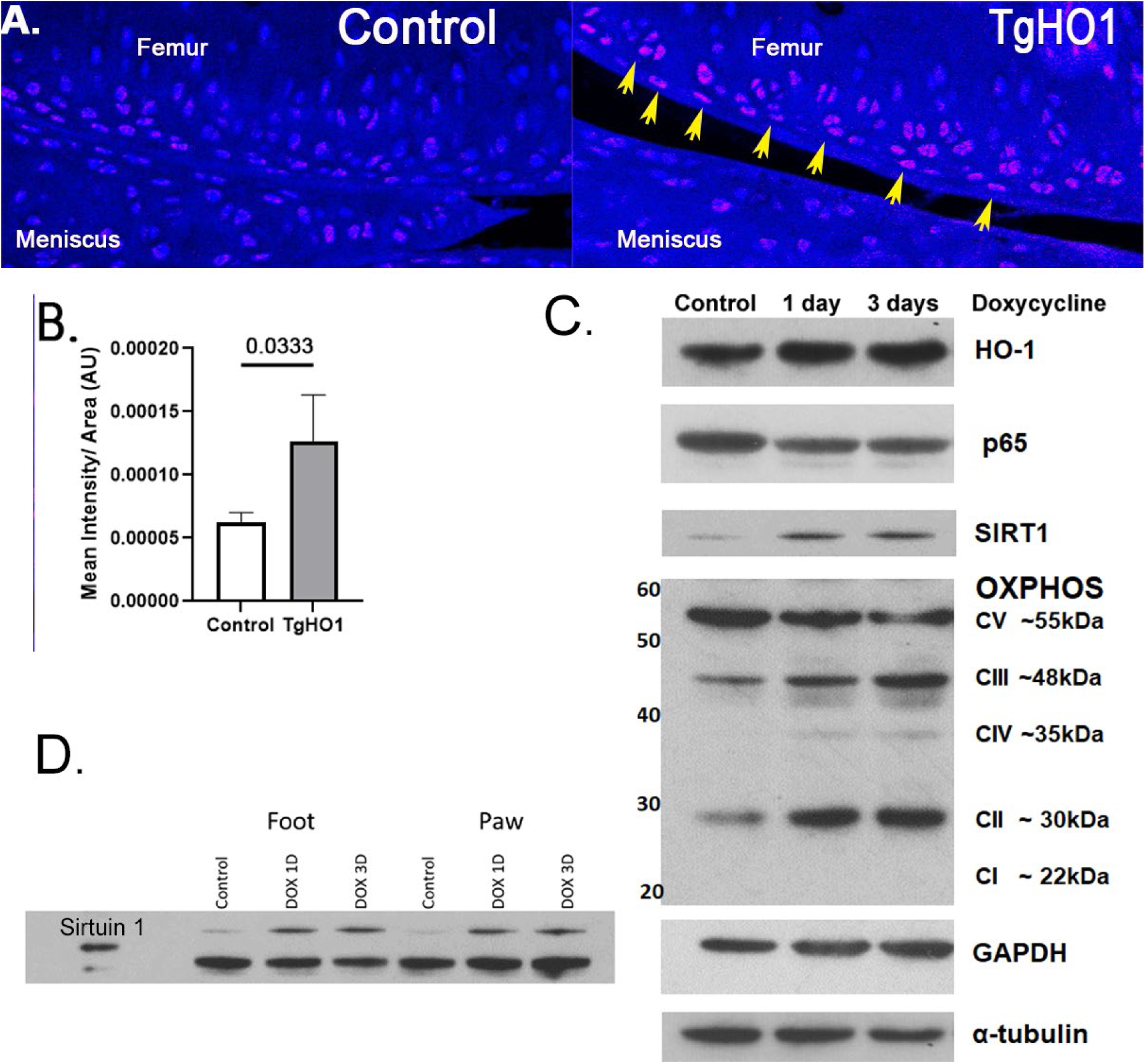
HO1 overexpression in articular chondrocytes increases mitochondrial metabolism and decreases pro-inflammatory signaling. Immunohistochemical staining of HO1 in articular cartilage shows increased staining in articular chondrocytes after 2 days of doxycycline feeding (A). These increases are not observed elsewhere in the joint. Western blotting of protein homogenates from the joints of these animals show induction of electron transport chain complexes II, III, and IV, suggesting increased respiratory capacity (B). We also note reduced total p65 (B). (C) Samples also showed increased Sirt1, supporting the hypothesis that mitochondrial biogenesis was stimulated. (D) Sirt1 western blotting shows induction of the protein both 1 and 3 days after initiation of doxycycline feeding.

### COF Augments Articular Chondrocyte GSH Metabolism and Protects Against Injury

Prior reports of antioxidant benefits from CO as well as the pro-mitochondrial effects of COF in cartilage suggested to us that COF might be protective against mitochondrial damage and dysfunction previously shown mediate PTOA at the earliest stages after injury. We hypothesized that COF could prevent GSH oxidation from a stylized model of traumatic injury and tested this hypothesis by applying COF for 1 h before a 2 J/cm^2^ impact injury known to oxidize GSH *in situ*. To provide detailed views of GSH status at the articular surface, we utilized the live cell stain MCB. This dye fluoresces blue in the presence of reduced GSH (vs oxidized) and this intensity decreases in the presence of decreased reduced GSH (i.e. dimmer blue = greater oxidation) [46, 47]. To demonstrate these specificities in articular cartilage we applied either buthionine sulfoximine (BSO) for 24 h before staining to inhibit glutathione synthesis or ethacrynic acid (ETA) immediately before staining to inhibit glutathione-S-transferases that contribute to MCB activation by GSH. Both inhibitors decreased MCB by greater than 90%, **Fig. 3A**.

**Fig. 3.**
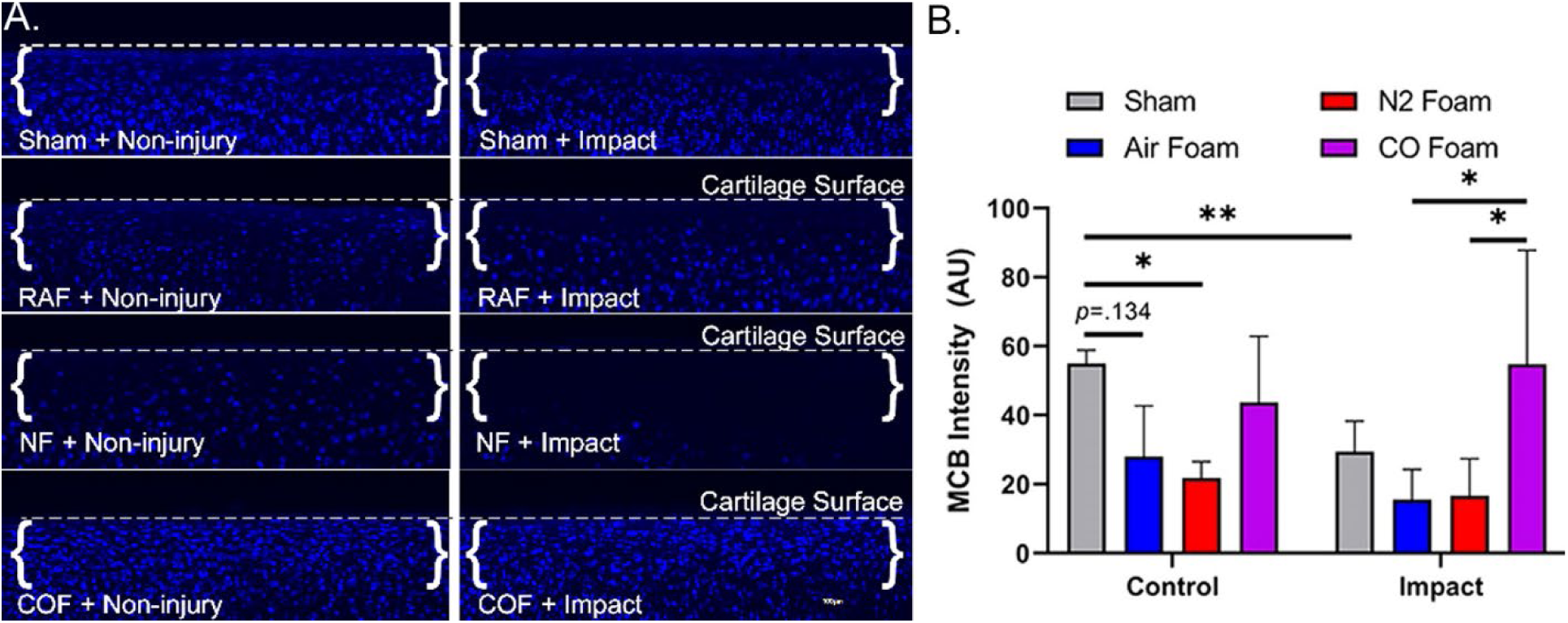
CO augments articular chondrocyte GSH metabolism to prevent damage. Osteochondral tissue was impacted with 2.0 J/cm2 impacts immediately after CO or control foam treatments. Control foams incurred losses in MCB that suggest redox injury (representative images shown in A, quantified in B). With impact, control foams did not prevent loss of MCB staining, while CO prevented oxidation of GSH indicated by this stain. n = 5, p < 0.05 by two-way ANOVA.

After exposure to either RAF, nitrogen foam (NF), or COF, explants were impacted, and returned to normal growth media for 24 h. MCB of cross sections of the articular surface showed that RAF and NF decreased MCB fluorescence intensity to a comparable degree to mechanical injury whereas COF maintained MCB staining in either setting, **Fig. 3B**. Oxidation in the absence of mechanical injury not observed in sham groups suggest that extended exposure to RAF and NF are injurious to a similar extent to mechanical injury, potentially because these foams are constructed with hypo-osmolar PBS relative to growth media [48]. We observed significant damage in the NF treated explants including some cell death, suggesting important distinctions between NF and COF despite both being constructed without oxygen gas. This demonstrates that CO’s specific biochemical activity is important to its benefits.

### Intraarticular CO is Non-Toxic and Improves Cartilage Health *In Situ*

Beyond the data above, COF provides ideal physical characteristics for intraarticular injection that also confer lubrication [16] and may prove especially useful by rapidly delivering CO to the entirety of the intraarticular joint. However, prior to clinical implementation, it is also crucial to demonstrate the safety and efficacy of this material. To first demonstrate that intraarticular COF rapidly filled the joint space after injection, we gave up to 4 ml injections of COF containing contrast agent to the medial compartments of the stifles of New Zealand White rabbits immediately post-mortem. This result was obtained with both rabbit fluoroscopic images and murine CT imaging after injection of the same contrast agent containing COF, **Fig. 4A and B**.

**Fig. 4.**
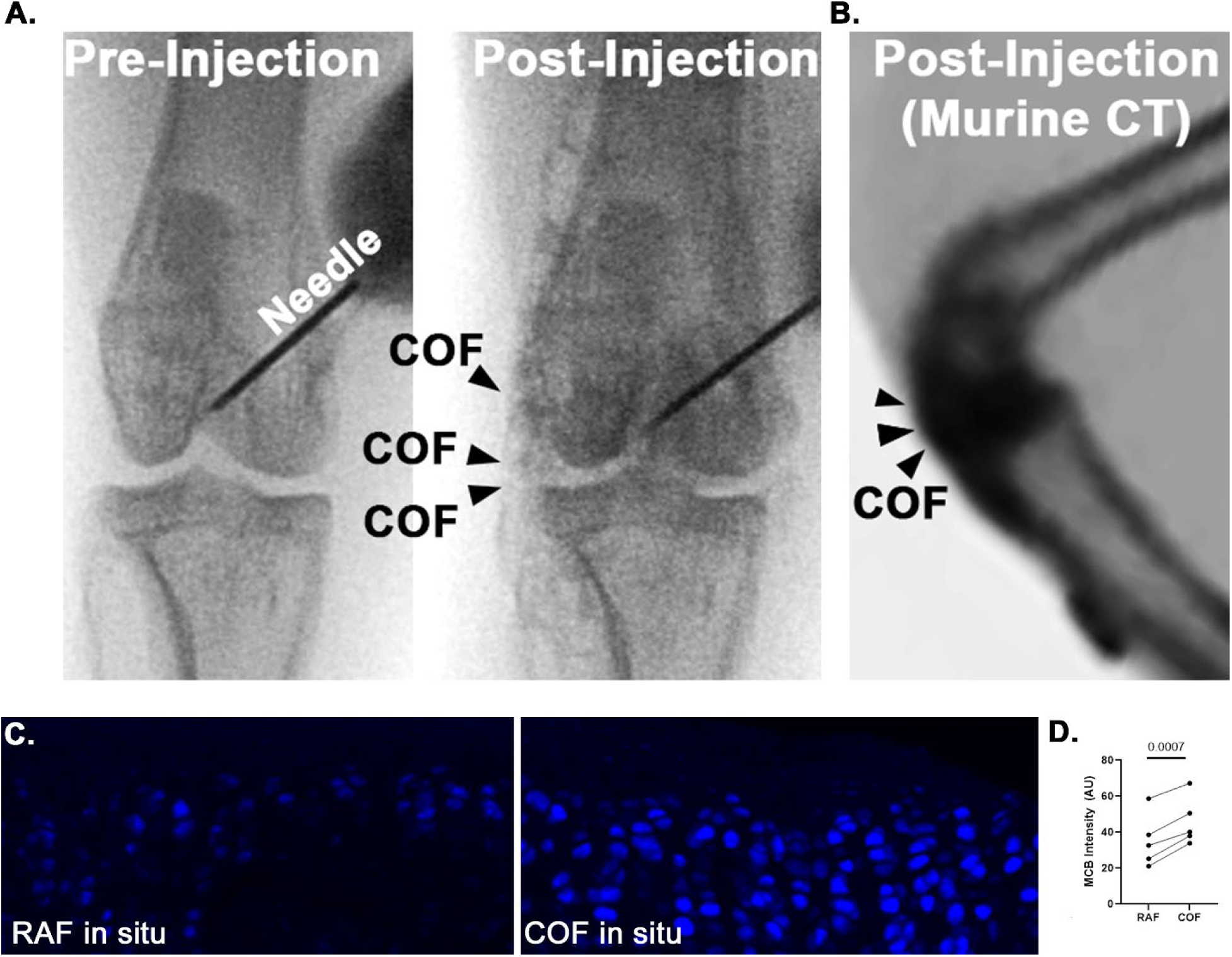
Intraarticular CO is non-toxic to chondrocytes and improves cartilage health in situ. Fluoroscopy of iodinated COF shows how intraarticular injection can fill the stifle joints of rabbits (A) and mice (B). Rabbit joints exposed to CO in this manner show increased MCB staining compared to contralateral joints exposed to RAF (shown in C and quantified in D). n = 5, p = 0.0007 by pairwise t-test for MCB analyses.

To interrogate whether COF has similar protective effects to our *in vitro* model, we utilized MCB staining of these rabbit stifles *ex vivo* 24 h after exposure. Rabbits were given RAF in one stifle and COF in the contralateral stifle, then 30 min later the limb was harvested, the articular surface was cut from the rest of the bone, and this was cultured for 24 h. This was intended to examine whether foam delivered while the joint was intact could provide similar significant increases in MCB intensity as was observed with osteochondral explants. We hypothesized that COF increased reduced GSH signal indicated by MCB *in situ*. COF treated cartilage showed significantly higher MCB fluorescent intensity than cartilage from pairwise contralateral joints exposed to comparable RAF doses, **Fig. 4C and D**.

To demonstrate the safety of this material, live rabbits were given either RAF or COF injections to the medial compartment of their left stifle. Animals were sacrificed either 1 week or 4 weeks after injection and articular surfaces of the tibia and femur were imaged for cell viability while synovia were fixed and prepared for histological staining with hematoxylin and eosin. This experiment demonstrated no indications of thickening or inflammation of the synovium in any of the rabbits given intraarticular injections of RAF or COF, **Fig. 5A-C**. We also observed no decreases in articular chondrocyte viability, **Fig. 5D**. These data demonstrated that 1 week after injection and 4 weeks after injection of COF the tissues of the joint appear completely normal, viable, and undamaged by the material. Because rabbit joints are too large for intact analyses, we injected RAF or COF in murine stifles and examined CO concentrations within the stifle 24 and 72 h later, seeing significant CO remaining in the joint 24 h after injection that does not remain 3 days after injection, **Fig. 5E**. This demonstrates that significant amounts of CO are retained for at least 24 h after injection.

**Fig. 5.**
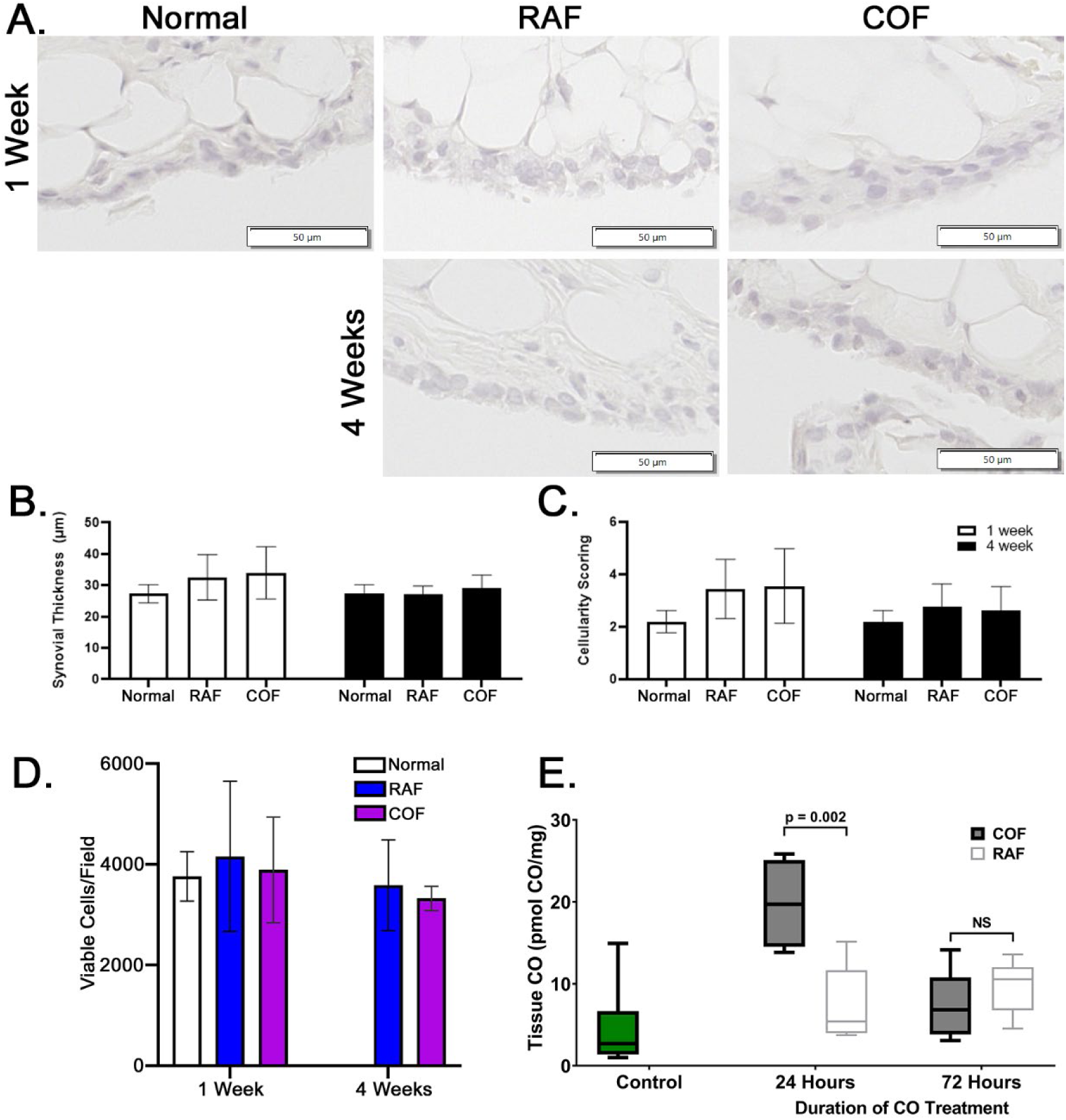
Intraarticular COF injection is Safe and Non-Toxic. (A) Representative images of synovia captured from rabbits either 1 week or 4 weeks after COF or RAF injection demonstrate no indications of thickening or increased infiltration by immune cells (quantified in B and C). No significant differences were observed, n = 10, analyzed via two-way ANOVA. Analyses of live cell densities from rabbits 1 week or 4 weeks after COF or RAF injection demonstrated no indications of any changes in live cell density at either time point using viable cell counts per field (D). Intact mouse stifle joints were analyzed for CO content 24 and 72 h after injection, n = 6 (E).

Lastly, to demonstrate proof-of-concept that HO1 expression in articular chondrocytes was sufficient to alter the course of PTOA after injury, we induced HO1 in double transgenic mice as shown in **Fig. 2** for the fourth week (days 22-28 after surgery) after meniscal destabilization procedures [49] and then animals were returned to normal chow at the start of week 5 until harvest 8 total weeks after meniscal destabilization. Average and maximum OARSI scoring by blinded, trained reviewers yielded p < 0.09 by two-way ANOVA followed by Kruskal Wallis non-parametric testing (n = 4), **Fig. 6**. These data support the hypothesis that inducing chondrocyte heme metabolism with HO1 can alter the course of PTOA after meniscal injury.

**Fig. 6.**
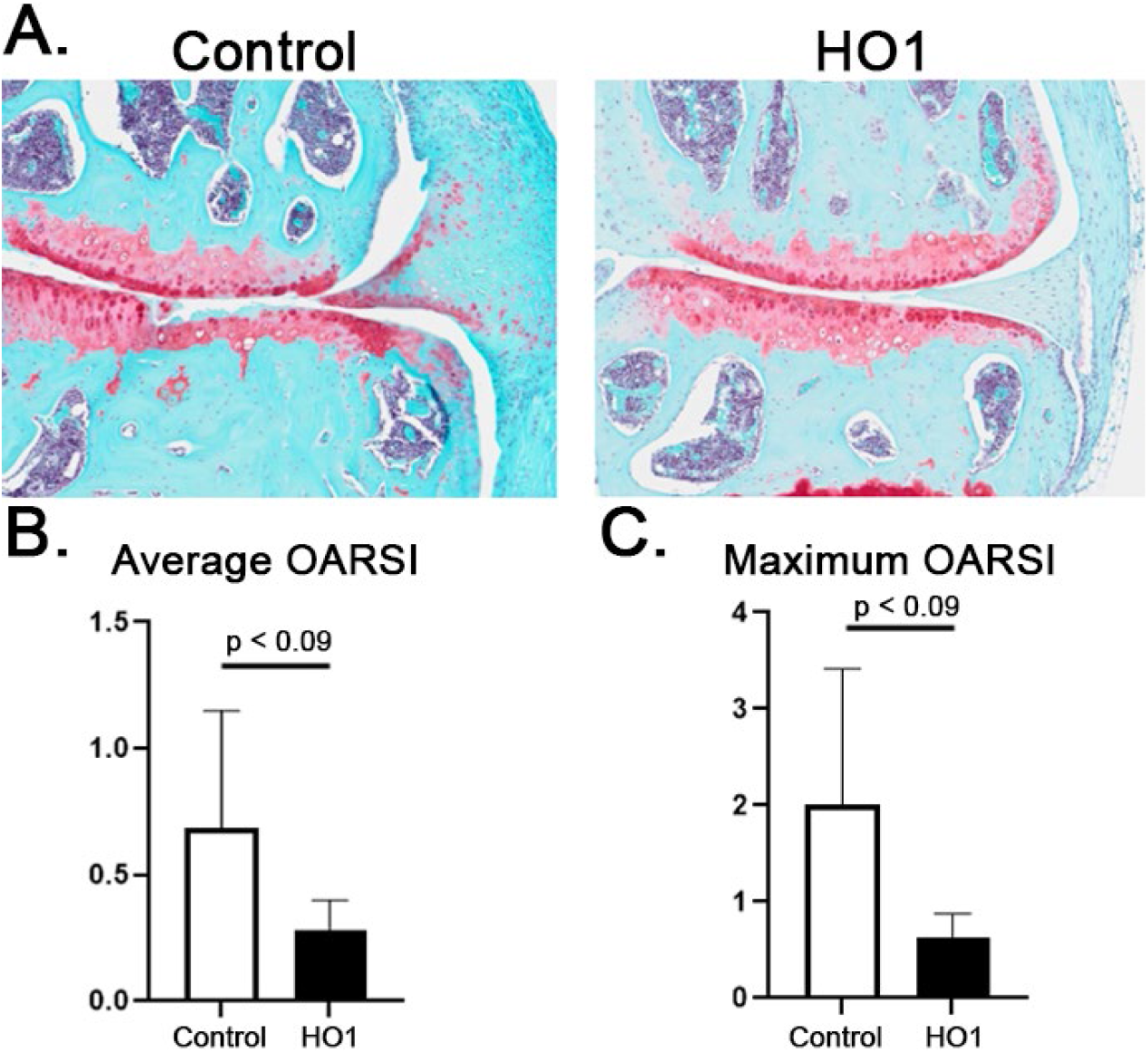
Activation of HO1 in Articular Chondrocytes Prevents PTOA. Mice given destabilization of the medial meniscus (DMM) reliably develop PTOA by 8-10 weeks. TgHO1 mice given doxycycline containing chow (625 mg/kg) for 1 week during the fourth week after injury show decreased average (B) and maximum (C) OARSI scores, suggesting HO1 prevents PTOA. N = 4, p = 0.0857 by Kruskal Wallis.

## [Discussion]

Articular cartilage injury contributes to PTOA *via* oxidation and mitochondrial dysfunction [25, 33, 50]. There are also mitochondrial abnormalities in chondrocytes later in disease progression [51]. This has led to enthusiasm in the orthopedic community for improving mitochondrial function and mitigating redox stress *via* “mitochondrial reprogramming”-based approaches that might prevent PTOA [25, 52]. Despite a history as a “silent killer”, scientific literature has shown therapeutic benefits of CO in line with these types of approaches [1-3]. We propose that activation of heme metabolism by CO causes prototypic mitochondrial reprogramming not only of great value to translational efforts, but to understanding cartilage healing processes as well. In this study, we have shown how this gasotransmitter provides an ideal adjuvant to intraarticular injuries that may be of value for patients with recently injured joints or joints that have begun to progress to PTOA. COF rapidly induced HO1 and increased healthy mitochondrial respiratory function in large animal tissue explants. COF also protected against GSH oxidation *in vitro* and *ex vivo* both in the presence and absence of mechanical injury without incurring any toxicity or inflammation on its own. Increases in mitochondrial respiration with HO1-induction and upregulation of heme metabolism in explant cartilage were reproduced *in vivo* using a cartilage-specific overexpressor of HO1. These data demonstrated acute, protective effects from COF to articular cartilage that may be efficacious for protecting against the stresses of intraarticular procedures when applied immediately pre-operatively.

We have included various strengths in the design of this study to increase the rigor and reproducibility of the articular cartilage biology under study. We included *in vitro* large animal explant models, *ex vivo* models of rabbit and mouse stifle joints, and *in vivo* animal models with consistent effects upon heme metabolism, mitochondria and GSH. The protective effects of CO were consistent across multiple species, with each demonstrating increased mitochondrial readouts as well as improved GSH status. We assessed GSH using multiple methods, each with detailed controls or standard curves as appropriate, to provide a more comprehensive understanding of the redox status of chondrocytes *in situ*. Similarly, we have adopted multiple approaches for investigating mitochondria, including both expression and activities of respiratory complexes as well as a variety of associated pro-mitochondrial pathways.

Several challenges and limitations to this study became clear during experimentation. It would be valuable to measure the oxygen consumption rate while CO is present and as CO is replaced with normal atmosphere; however, given the technical details of the mitochondrial stress test setup, it is unclear whether the foam can be rinsed effectively from extracellular flux analyzer wells to provide reliable, immediate quantitation. In lieu of this, we have chosen to manually confirm foam removal and then initiate mitochondrial stress tests, yielding the data provided. We note that the paradox that a molecule known to inhibit respiration would augment mitochondrial activity after removal has been observed in other tissues [53, 54]. More detailed, depth- and diffusion-focused research could provide interesting kinetic views of how CO’s chemical activity alters short-term chondrocyte metabolic behavior. Because the delivery of CO from COF relies on diffusion, depth- and time-specific effects may be important to future preclinical and translational efforts. Large animal model systems may prove especially useful in future studies describing time-dependent and dose-dependent features of gasosignaling intraarticular biology.

We have already noted the paradox that CO inhibits heme-containing proteins, including the mitochondrial complexes, yet we observed an increase in mitochondrial respiration and expression with CO or HO1 overexpression. This was observed as soon as 1 h after COF and persisted to 24 h after administration. Increases in reduced GSH also persisted to 24 h. In this way, we suggest that activation of heme metabolism in articular chondrocytes induces endogenous mitochondrial reprogramming that can provide lasting benefits to articular cartilage. By stimulating mitochondrial biogenesis, supporting antioxidant activities through GSH, and thereby coordinating a return to healthy oxygen metabolism after exposures to COF, this material might be applied immediately before any procedure where low osmolarity, low temperature, high oxygen saline, and the other rigors of intraarticular surgery might present a threat to cartilage health. The direct effects of COF (healthier, more functional mitochondria and robust antioxidant defenses) and, by extension, the activation of heme metabolism are ideally suited to this setting.

This study has concentrated on specific protective effects of COF to bolster application of COF within the context of preventive care for recently injured but otherwise healthy cartilage; however, we anticipate similar benefits in the other tissues of the articular joint or for cartilage already along the continuum of PTOA and studies are currently underway to delineate these benefits. Long term, we expect this material and local activation of heme metabolism to protect against development of PTOA not only by coordinating chondrocyte intracellular metabolism, but also by coordinating metabolic function and related pro-inflammatory/anti-inflammatory signaling throughout the joint. Coordinated protection and restoration of cartilage metabolism coincident with stimulation of the synovium by COF, specifically upregulation of anti-inflammatory macrophages, may represent not only an ideal system in which to examine intraarticular gasosignaling and cartilage-synovium crosstalk, but may also provide revolutionary therapeutic benefits. The COF shown here providing these benefits consists of simple, GRAS materials which confer meaningful, lasting safety and biocompatibility. COF is easily injectable into rodent joints, thus represents a realistic solution for human intraarticular injection. Thus, COF is an ideal adjuvant to orthopedic surgical care in any context where damage to the cartilage is a concern.

## [ACKNOWLEDGMENTS]

We are grateful to be able to conduct this study with supports from NIAMS K99/R00 AR070914, DOD #W81XWH-11-1-0583, and DOD #W81XWH-10-1-0864 grant, NCI K08CA276908, V Scholars Award, National Cancer Institute Core Grant (P30 CA086862), University of Iowa Holden Comprehensive Cancer Center, ACS-IRG-21-141-46, National Science Foundation (# 1927616), University of Iowa Hospitals and Clinics Start-up, Department of Radiation Oncology and Department of Orthopedics and Rehabilitation of The University of Iowa, and Orthopedic Histopathology Service of The University of Iowa.

## [AUTHOR CONTRIBUTIONS]

JDB, SL, PCGC, and LDM assisted in foam mixing and administration. SL, PCGC, MRH, JSF, KJL, LDM, MYS, EW participated in bovine, lapine, and murine experimentation. ABP and MST conducted CO concentration measurements. SL, PCGC, KJL, and BAW assisted in extracellular flux analyses and biochemical analyses. SL, JSF, KJL, and MCC assisted in the analysis of all data as well as interpretation. SL and MCC wrote the manuscript. All authors participated in editing the manuscript.

## Figures

**Supplemental Fig. 1.**
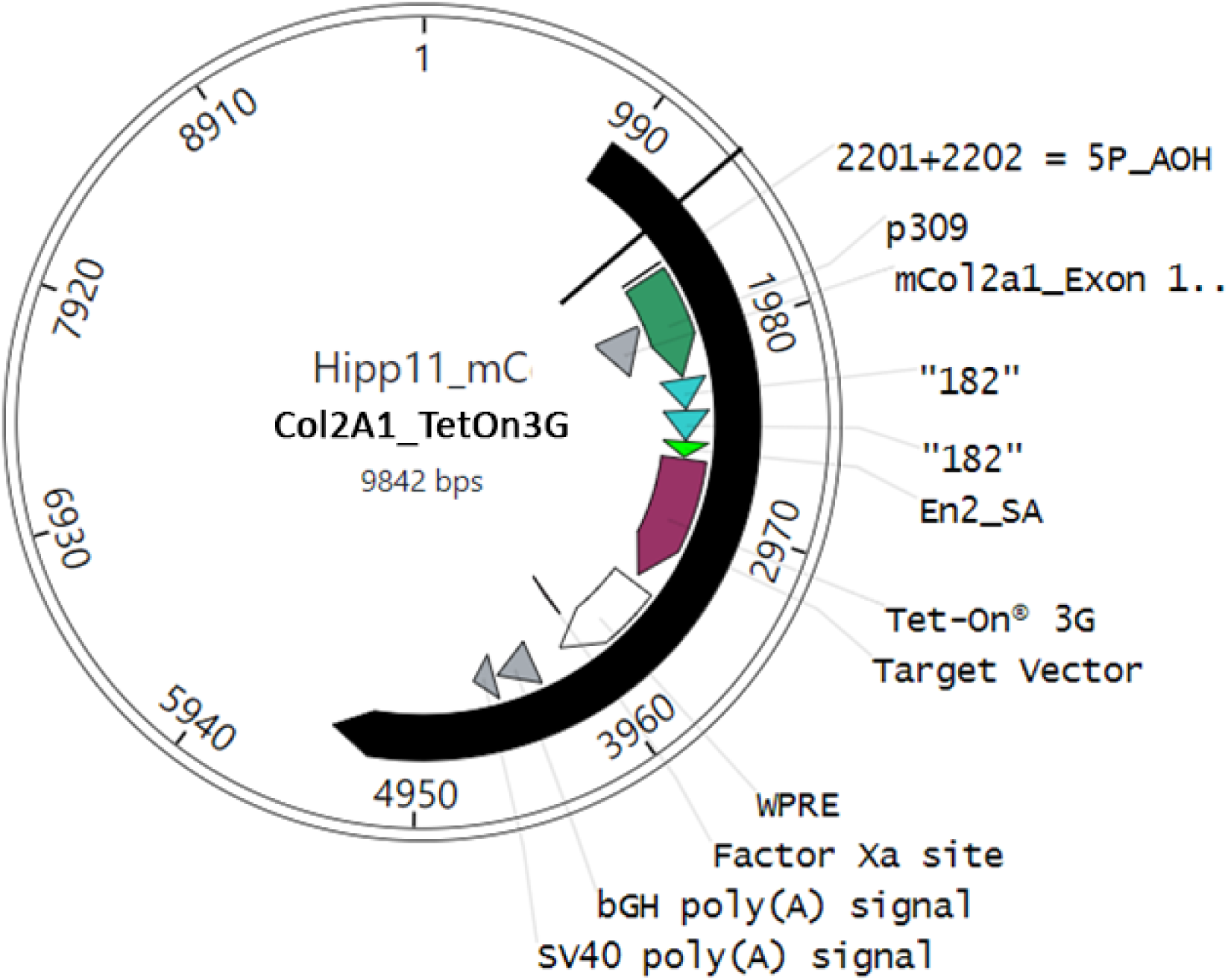
Transgene Inserted for Creation of Col2A1-TetOn Mice. the inserted Col2A1-TetOn transgene (black) contains the necessary Hip11 (H11) sequences for insertion, an SV40 polyadenylation signal, the previously characterized 2x182 bp doublet taken from the Col2A1 promoter sequence, a minimal woodchuck hepatitis virus post-transcriptional regulatory element (WPRE), the TetOn 3G gene, a Factor Xa site and polyadenylation signal, and the second H11 sequence.

**Supplemental Fig. 2.**
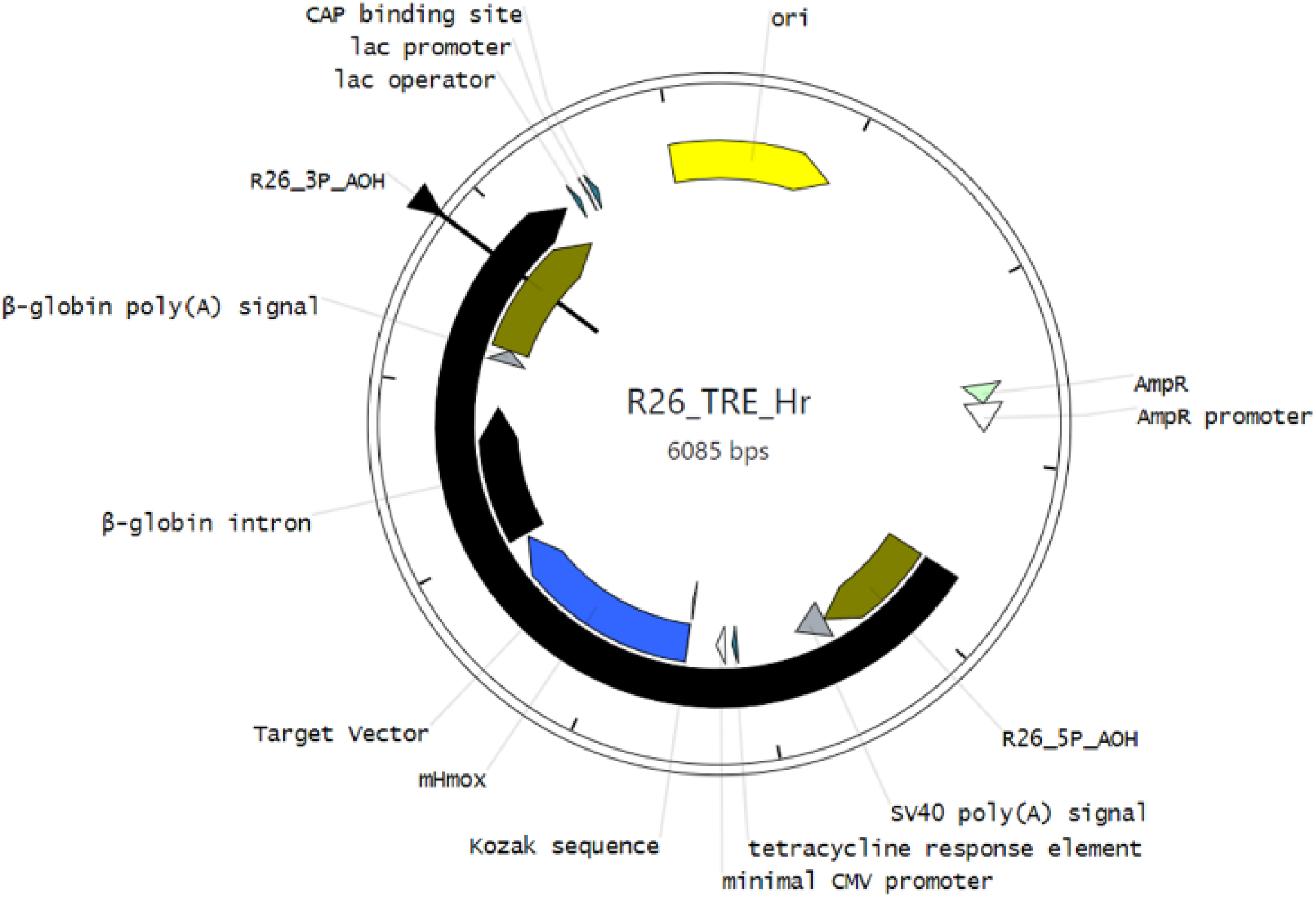
Transgene Inserted for Creation of TRE-mHMOX1 Mice. The inserted TRE-mHMOX1 transgene (black) contains the necessary R26 sequences for insertion, an SV40 polyadenylation signal, the tet response element critical for rtTA*M2 binding (7 repeats of TCCCTATCAGTGATAGAGA separated by spacer sequences), a minimal cytomegalovirus promoter, the Kozak sequence and mHMOX1 gene, a β-globin intron and polyadenylation signal, and the second R26 sequence.

## References

1. Lee, P.J., et al., Regulation of heme oxygenase-1 expression in vivo and in vitro in hyperoxic lung injury. American journal of respiratory cell and molecular biology, 1996. 14(6): p. 556–568.

2. Zhang, X., et al., Mitogen-activated protein kinases regulate HO-1 gene transcription after ischemia-reperfusion lung injury. American Journal of Physiology-Lung Cellular and Molecular Physiology, 2002. 283(4): p. L815–L829.

3. W Ryter, S. and A. MK Choi, Heme oxygenase-1/carbon monoxide: novel therapeutic strategies in critical care medicine. Current drug targets, 2010. 11(12): p. 1485–1494.

4. Haider, A., et al., Regulation of cyclooxygenase by the heme-heme oxygenase system in microvessel endothelial cells. Journal of Pharmacology and Experimental Therapeutics, 2002. 300(1): p. 188–194.

5. Quan, S., et al., Functional expression of human heme oxygenase-1 (HO-1) driven by HO-1 promoter in vitro and in vivo. Journal of cellular biochemistry, 2002. 85(2): p. 410–421.

6. Converso, D.P., et al. HO-1 is located in liver mitochondria and modulates mitochondrial heme content and metabolism. 2006. Federation of American Societies for Experimental Biology.

7. Bansal, S., G. Biswas, and N.G. Avadhani, Mitochondria-targeted heme oxygenase-1 induces oxidative stress and mitochondrial dysfunction in macrophages, kidney fibroblasts and in chronic alcohol hepatotoxicity. Redox Biol, 2014. 2: p. 273–83.

8. Motterlini, R. and L.E. Otterbein, The therapeutic potential of carbon monoxide. Nat Rev Drug Discov, 2010. 9(9): p. 728–43.

9. Motterlini, R., B.E. Mann, and R. Foresti, Therapeutic applications of carbon monoxide-releasing molecules. Expert Opin Investig Drugs, 2005. 14(11): p. 1305–18.

10. Motterlini, R. and R. Foresti, Carbon-Monoxide-Releasing Molecules (CO-RMs): A Stratagem to Emulate the Beneficial Effects of Heme Oxygenase-1, in Heme Oxygenase: The Elegant Orchestration of Its Products in Medicine, L.E. Otterbein and B.S. Zuckerbraun, Editors. 2005, Nova Biomedical Books: New York. p. 191-210.

11. Benallaoua, M., et al., Pharmacologic induction of heme oxygenase 1 reduces acute inflammatory arthritis in mice. Arthritis Rheum, 2007. 56(8): p. 2585–94.

12. Guillén, M., et al., Haem oxygenase-1 regulates catabolic and anabolic processes in osteoarthritic chondrocytes. J Pathol, 2008. 214(4): p. 515–22.

13. Bonelli, M., et al., Heme oxygenase-1 end-products carbon monoxide and biliverdin ameliorate murine collagen induced arthritis. Clin Exp Rheumatol, 2012. 30(1): p. 73–8.

14. Silva, G., et al., Heme Oxygenase-1: A Protective Gene that Regulates Inflmation and Immunity, in Heme Oxygenase: The Elegant Orchestration of Its Products in Medicine, L.E. Otterbein and B.S. Zuckerbraun, Editors. 2005, Nova Biomedical Books: New York. p. 141-169.

15. Sharma, H.S., et al., Coordinated expression of heme oxygenase-1 and ubiquitin in the porcine heart subjected to ischemia and reperfusion. Mol Cell Biochem, 1996. 157(1-2): p. 111–6.

16. Byrne, J.D., et al., Delivery of therapeutic carbon monoxide by gas-entrapping materials. Sci Transl Med, 2022. 14(651): p. eabl4135.

17. Belcher, J.D. and G.M. Vercellotti, Heme Oxygenase-1: A Potential Modulator of Inflammation and Vasoocclution in Sickle Cell Disease, in Heme Oxygenase: The Elegant Orchestration of Its Products in Medicine, L.E. Otterbein and B.S. Zuckerbraun, Editors. 2005, Nova Biomedical Books: New York. p. 97-111.

18. Ryter, S.W., D. Morse, and A.M. Choi, Carbon monoxide and bilirubin: potential therapies for pulmonary/vascular injury and disease. Am J Respir Cell Mol Biol, 2007. 36(2): p. 175–82.

19. Converso, D.P., et al., HO-1 is located in liver mitochondria and modulates mitochondrial heme content and metabolism. FASEB J, 2006. 20(8): p. 1236–8.

20. Blanco, F.J., M.J. López-Armada, and E. Maneiro, Mitochondrial dysfunction in osteoarthritis. Mitochondrion, 2004. 4(5-6): p. 715–28.

21. Johnson, K., et al., Mitochondrial oxidative phosphorylation is a downstream regulator of nitric oxide effects on chondrocyte matrix synthesis and mineralization. Arthritis Rheum, 2000. 43(7): p. 1560–70.

22. Ramakrishnan, P., et al., Oxidant conditioning protects cartilage from mechanically induced damage. J Orthop Res, 2010. 28(7): p. 914–20.

23. Brandl, A., et al., Oxidative stress induces senescence in chondrocytes. J Orthop Res, 2011. 29(7): p. 1114–20.

24. Coleman, M.C., et al., Differential Effects of Superoxide Dismutase Mimetics after Mechanical Overload of Articular Cartilage. Antioxidants (Basel), 2017. 6(4).

25. Coleman, M.C., et al., Targeting mitochondrial responses to intra-articular fracture to prevent posttraumatic osteoarthritis. Sci Transl Med, 2018. 10(427).

26. Blanco, F.J., I. Rego, and C. Ruiz-Romero, The role of mitochondria in osteoarthritis. Nat Rev Rheumatol, 2011. 7(3): p. 161–9.

27. Bauer, M.S., J.C. Woodard, and J.P. Weigel, Effects of exposure to ambient air on articular cartilage of rabbits. Am J Vet Res, 1986. 47(6): p. 1268–70.

28. Zhou, S., Z. Cui, and J.P. Urban, Factors influencing the oxygen concentration gradient from the synovial surface of articular cartilage to the cartilage-bone interface: a modeling study. Arthritis Rheum, 2004. 50(12): p. 3915–24.

29. Mignotte, F., et al., Mitochondrial biogenesis in rabbit articular chondrocytes transferred to culture. Biol Cell, 1991. 71(1-2): p. 67–72.

30. Hwang, J.W., et al., Effects of solvent osmolarity and viscosity on cartilage energy dissipation under high-frequency loading. J Mech Behav Biomed Mater, 2022. 126: p. 105014.

31. Villanueva, I., N.L. Bishop, and S.J. Bryant, Medium osmolarity and pericellular matrix development improves chondrocyte survival when photoencapsulated in poly(ethylene glycol) hydrogels at low densities. Tissue Eng Part A, 2009. 15(10): p. 3037–48.

32. Loret, B. and F.M. Simões, Effects of pH on transport properties of articular cartilages. Biomech Model Mechanobiol, 2010. 9(1): p. 45–63.

33. Coleman, M.C., et al., Injurious Loading of Articular Cartilage Compromises Chondrocyte Respiratory Function. Arthritis Rheumatol, 2016. 68(3): p. 662–71.

34. Birch-Machin, M.A., et al., An evaluation of the measurement of the activities of complexes I-IV in the respiratory chain of human skeletal muscle mitochondria. Biochem Med Metab Biol, 1994. 51(1): p. 35–42.

35. Gossen M, Freundlieb S, Bender G, Müller G, Hillen W, Bujard H. Transcriptional activation by tetracyclines in mammalian cells. Science. 1995;268(5218):1766-1769.

36. Zhou G, Garofalo S, Mukhopadhyay K, et al. A 182 bp fragment of the mouse pro alpha 1(II) collagen gene is sufficient to direct chondrocyte expression in transgenic mice. J Cell Sci. 1995;108 (Pt 12):3677–3684.

37. Glasson SS, Blanchet TJ, Morris EA. The surgical destabilization of the medial meniscus (DMM) model of osteoarthritis in the 129/SvEv mouse. Osteoarthritis Cartilage. 2007 Sep;15(9):1061–9.

38. Hines, M.R., et al., Extracellular biomolecular free radical formation during injury. Free Radic Biol Med, 2022. 188: p. 175–184.

39. Yang, L., et al., Deep Learning for Chondrocyte Identification in Automated Histological Analysis of Articular Cartilage. Iowa Orthop J, 2019. 39(2): p. 1–8.

40. Griffith, O.W., Determination of glutathione and glutathione disulfide using glutathione reductase and 2-vinylpyridine. Anal Biochem, 1980. 106(1): p. 207–12.

41. Zuckerbraun, B.S., et al., Carbon monoxide protects against liver failure through nitric oxide-induced heme oxygenase 1. J Exp Med, 2003. 198(11): p. 1707–16.

42. Nakao, A., et al., Protective effect of carbon monoxide inhalation for cold-preserved small intestinal grafts. Surgery, 2003. 134(2): p. 285–92.

43. Ghanta, S., et al., Mesenchymal Stromal Cells Deficient in Autophagy Proteins Are Susceptible to Oxidative Injury and Mitochondrial Dysfunction. Am J Respir Cell Mol Biol, 2017. 56(3): p. 300–309.

44. Lancel, S., et al., Carbon monoxide rescues mice from lethal sepsis by supporting mitochondrial energetic metabolism and activating mitochondrial biogenesis. J Pharmacol Exp Ther, 2009. 329(2): p. 641–8.

45. Piantadosi, C.A., et al., Heme oxygenase-1 regulates cardiac mitochondrial biogenesis via Nrf2-mediated transcriptional control of nuclear respiratory factor-1. Circ Res, 2008. 103(11): p. 1232–40.

46. Rice, G.C., et al., Quantitative analysis of cellular glutathione by flow cytometry utilizing monochlorobimane: some applications to radiation and drug resistance in vitro and in vivo. Cancer Res, 1986. 46(12 Pt 1): p. 6105-10.

47. Cook, J.A., et al., Use of monochlorobimane for glutathione measurements in hamster and human tumor cell lines. Int J Radiat Oncol Biol Phys, 1989. 16(5): p. 1321–4.

48. Ozturk, S.S. and B.O. Palsson, Effect of medium osmolarity on hybridoma growth, metabolism, and antibody production. Biotechnology and bioengineering, 1991. 37(10): p. 989–993.

49. Glasson SS, Blanchet TJ, Morris EA. The surgical destabilization of the medial meniscus (DMM) model of osteoarthritis in the 129/SvEv mouse. Osteoarthritis Cartilage. 2007 Sep;15(9):1061–9.

50. Gavriilidis, C., et al., Mitochondrial dysfunction in osteoarthritis is associated with down-regulation of superoxide dismutase 2. Arthritis Rheum, 2013. 65(2): p. 378–87.

51. Buckwalter, J.A., et al., The Roles of Mechanical Stresses in the Pathogenesis of Osteoarthritis: Implications for Treatment of Joint Injuries. Cartilage, 2013. 4(4): p. 286–294.

52. Ansari, M.Y., et al., Parkin clearance of dysfunctional mitochondria regulates ROS levels and increases survival of human chondrocytes. Osteoarthritis Cartilage, 2018. 26(8): p. 1087–1097.

53. Wang, L., et al., Inhibition of mitochondrial respiration under hypoxia and increased antioxidant activity after reoxygenation of Tribolium castaneum. PLoS One, 2018. 13(6): p. e0199056.

54. Morse, P.T., et al., Phosphorylations and Acetylations of Cytochrome c Control Mitochondrial Respiration, Mitochondrial Membrane Potential, Energy, ROS, and Apoptosis. Cells, 2024. 13(6).

